# Maternal behavior in Sumatran orangutans (*Pongo abelii*) is modulated by mother-offspring characteristics and socioecological factors

**DOI:** 10.1101/2024.03.02.582963

**Authors:** T Revathe, Roger Mundry, Sri Suci Utami-Atmoko, Paul-Christian Bürkner, Maria A. van Noordwijk, Caroline Schuppli

**Author notes:** Corresponding author: Correspondence to T Revathe, Development and Evolution of Cognition Group Max Planck Institute of Animal Behavior Bücklestraße 5a, 78467 Konstanz Germany.

## Abstract

Mammalian mothers flexibly invest in their offspring to maximize their lifetime fitness. Flexible maternal investment may be particularly important in large-brained species with prolonged maternal care – e.g., in great apes. We investigated the effects of socioecological factors and mother-offspring characteristics on nine maternal behaviors in wild Sumatran orangutans (*Pongo abelii*; *N*=22 mother-offspring pairs; over 11,200 hours of focal data; collected from 2007-2022) using generalized linear mixed models. The behaviors fall under four maternal functions: locomotory support (carrying), skill acquisition support (feeding in proximity, food transfer), protective proximity maintenance (body contact and proximity initiation, following), and independence promotion (body contact and proximity termination, avoiding). Mother’s parity was not significantly associated with any maternal behavior. Mothers were more likely to show locomotory support, skill acquisition support, and protective proximity maintenance toward younger than older offspring; whereas they were more likely to promote independence in older than younger offspring. Mothers with male offspring were more likely to show skill acquisition support to their offspring than those with female offspring. With increasing food availability, skill acquisition support reduced. With increasing association size, mothers were more likely to show protective proximity maintenance and less likely to promote independence. When males were present, mothers were more likely to show locomotory support to their offspring. Sumatran orangutan mothers thus flexibly adjust offspring directed behavior in response to prevailing socioecological factors and mother-offspring characteristics. Our findings add support to the evolutionary theory that mammalian mothers flexibly invest in their offspring.

## INTRODUCTION

The mother-offspring relationship is of particular significance in the life of young mammals. This is especially true of large-brained altricial mammals with complex behavioral repertoires – such as cetaceans, elephants, and human and nonhuman primates, in which physical and behavioral development occurs over an extended postnatal period (Lee and Moss, 2011; Mann, 2019; Revathe et al., 2020; van Noordwijk & van Schaik, 2005). In such species, the developmental period stretches over multiple years, which means that individuals are likely to be confronted with numerous changes in their ecological and social environments. In most mammals, mothers bear the major or the sole responsibility of postnatal care (Clutton-Brock, 1991; Maestripieri, 2018; Nicolson, 1991; van Noordwijk, 2012). Mothers provide support for nutrition (through milk and sharing solid food: Brown et al., 2004; Hinde & Milligan, 2011; Johnson et al., 2010; Lee and Moss, 2011; Nair, 1989), and thermoregulation (through maintaining body contact: Lubach et al., 1992; Taber & Thomas, 1982). Furthermore, mothers actively maintain the distance between themselves and their dependents (Förster & Cords, 2002; Szabo & Duffus, 2008); facilitate their dependents’ independence acquisition (through breaking body contact: Maestripieri, 1995) and help in the formation of their social relationships (by choosing association partners and maintaining relationships with them: Fairbanks and McGuire, 1986; Murray et al., 2014). Mothers, along with other social unit members, protect their dependents against predators (Lee and Moss, 2011; Mann, 2019; Nicolson, 1991). In primates, mothers additionally provide locomotory support (through carrying and, in certain primates, bridging to cross canopy: Chappell et al., 2015; van Noordwijk & van Schaik, 2005), and serve as role models for social learning of skill and knowledge repertoires (Inoue-Nakamura & Matsuzawa, 1997; Jaeggi et al., 2008; Lonsdorf, 2006; Schuppli et al., 2016). Consequently, in the absence of mothers and their care, offspring mortality risk increases in mammals (e.g., Lahdenperä et al., 2016; Zipple et al., 2021).

Maternal investment is defined in terms of future reproductive costs incurred by the mother for reproductive effort allocated to her current offspring (Trivers, 1972). As maternal investment – e.g., gestation, lactation, and carrying – is time-consuming and energy-intensive, females are expected to balance the benefits and costs of investing in each of their offspring so as to maximize their lifetime reproductive success (Clutton-Brock, 1991; Lee et al., 1991). In several species, mothers flexibly adjust the quantity and quality of their investment in response to their and their offspring’s characteristics (e.g., parity, maternal age, offspring age and sex), as well as to their immediate ecological and social conditions (e.g., social unit size and composition), which are studied in detail in many primate species (Lee, 1987; Nicolson, 1987, 1991; van Noordwijk, 2012).

In primates, maternal age and parity often affect maternal investment, wherein offspring of younger/primiparous females have a lower body mass, face more neglect and abandonment, and consequently have reduced survival compared to those of older/multiparous females (Altmann and Alberts, 2005; Arlet et al., 2014; Fairbanks, 1996; but see Silk, 1991). Possible explanations include that younger/primiparous mothers face a trade-off between investing in their own growth vs. in their offspring’s (Setchell et al., 2002), therefore they may produce less milk than older/multiparous mothers (Hinde et al., 2009) or are less experienced and thus less skilled in rearing their offspring than older/multiparous mothers (Mitchell & Stevens, 1968).

In general, maternal investment decreases with increasing offspring age due to the offspring’s increasing ecological and social competence (Dias et al., 2018; Fairbanks, 1996; Jaeggi et al., 2008; Li et al., 2013; Mikeliban et al., 2021; Nishida and Turner, 1996). However, mothers and offspring may disagree over the period of investment due to an underlying conflict (parent-offspring conflict theory; Trivers, 1974). Concerning offspring sex, the Trivers-Willard hypothesis posits that in species in which body condition differentially affects the reproductive success of the sexes, mothers should bias their investment in favor of the sex with the potentially higher reproductive return when controlled for investment (males in polygynous species), provided mothers’ investment during dependency translates into offspring’s body condition in adulthood (Trivers & Willard, 1973). Studies on primates found inconsistent results for sex-biased maternal investment (Dettmer et al., 2016; Fairbanks, 1996; Leigh & Shea, 1995; Setchell et al., 2002; Silk, 1991; Tanaka, 1989).

Maternal duration of investment is hypothesized to have an inverse U-shaped relationship with maternal body condition in response to food availability, with mothers in average body condition investing more than mothers in poor or good body condition (Fairbanks & McGuire, 1995; Lee et al., 1991), especially in annual breeders. The quadratic relationship is explained with the rationale that mothers living in relatively poor-quality habitats may not be able to satisfy both their own and their offspring’s needs, and thus may reduce the period of investment (such as through early weaning age; Lee et al., 1991) to support their future reproduction at the expense of current offspring’s survival. Whereas mothers living in high-quality habitats are likely able to support their and their offspring’s needs, but can reduce the period of investment in the current offspring to support one’s future reproduction without increasing the risk of offspring mortality (Hauser & Fairbanks, 1988).

Mothers may also adjust their investment in response to the social risk posed by sociodemographic factors such as social unit size and composition along with dominance structure (Maestripieri, 1994; Nicolson, 1991). For example, it is hypothesized that in species with frequent aggressive interactions among social unit members, with an increase in social unit size mothers may invest in staying close to their offspring because of the perceived social risk to offspring (Maestripieri, 1994). When there is a risk of infanticide by males, mothers invest more in their offspring by staying closer to them in the presence compared to in the absence of particular males in the social unit (Fairbanks & McGuire, 1987; Scott et al., 2019, 2023).

The effects of mother-offspring characteristics, ecological, and sociodemographic variables on maternal investment have been studied in various terrestrial primates (e.g., vervet monkeys (*Cercopithecus aethiops sabaeus*): Fairbanks and McGuire, 1995; rhesus macaques (*Macaca mulatta*): Maestripieri, 2001; baboons (*Papio cynocephalus*): Nguyen et al., 2012; mandrills (*Mandrillus sphinx*): Setchell et al., 2002). However, most of these studies have looked at one or few factors influencing one or few forms of maternal investment rather than comprehensively investigating the effects of several factors on multiple forms of investment. Simultaneously testing for the effects of multiple known predictors of investment is important, as effects of certain predictors may only become apparent when controlling for the effects of other predictors (e.g., controlling for offspring age). Furthermore, studies on maternal investment in arboreal primates are comparatively rare (but see Arlet et al., 2014; Dias et al., 2018; Mikeliban et al., 2021; van Noordwijk & van Schaik, 2005). As arboreal primates face different ecological challenges compared to terrestrial primates (e.g., learning to cross gaps in the canopy without risking a fall), some arboreal primates show specialized maternal behaviors like bridging. Furthermore, the arboreal lifestyle entails high energetic costs, especially for large primates (e.g., climbing costs: Hanna & Schmitt, 2011; Thorpe and Crompton 2009), which may lead to constraints on maternal investment. Thus, the obvious differences in habitat, predation risk, diet composition, energetic constraints, and social characteristics between terrestrial and arboreal primates must be taken into account to understand the variation found in their patterns of maternal care.

Among arboreal primates, orangutans (*Pongo* spp.) are particularly interesting to study maternal care because they have the slowest life history, including the longest dependency period and inter-birth intervals of all nonhuman primates (van Adrichem et al., 2006; van Noordwijk et al., 2018). Females give birth to a single offspring at a time and provide care for 6-8 years of age (van Noordwijk et al., 2009). Mothers invest heavily in each offspring through prolonged nursing, carrying, and nest sharing (van Noordwijk & van Schaik, 2005). Furthermore, the orangutans’ predominantly arboreal lifestyle requires maternal care in the form of carrying and bridging to cross gaps in the canopy. Males disperse from their natal area and do not provide paternal care (Arora et al., 2012; Morrogh-Bernard et al., 2011; Nietlisbach et al., 2012; van Noordwijk et al., 2023). This, along with a prolonged period of lactation relative to gestation length (van Schaik, 2000), sexual size dimorphism (Utami Atmoko et al., 2009), and potential for contest competition between males for mating opportunities (Kunz et al., 2023; Spillmann et al., 2017; van Schaik & van Hooff, 1996), puts orangutans at a high risk of infanticide by males, even though there are no confirmed cases (Beaudrot et al., 2009; Knott et al., 2019; van Schaik & Janson, 2000).

Most orangutan habitats are characterized by low and fluctuating food availability (Marshall et al., 2009). Even though fruits are the orangutans’ main energy source, individuals survive on fallback foods during periods of low fruit availability (Morrogh-Bernard et al., 2009). Furthermore, the orangutans’ arboreal lifestyle and large body and brain size result in high energetic requirement. As a likely response to the energetic constraints, orangutans have developed behavioral and physiological adaptations to deal with periods of low food availability (e.g., reduced movement when food is scarce and exceptionally low basal metabolic rate (BMR); Morrogh-Bernard et al., 2009; Pontzer et al., 2010; Vogel et al., 2017).

As a result of low forest productivity and energetic constraints, associations are costly to maintain for orangutans (Kunz et al., 2021; van Schaik, 1999). They are generally semi-solitary (Setia et al., 2009), but mothers with dependent offspring spend up to 50% of their time in association with others in some sites (depending on population; Roth et al., 2020; van Noordwijk et al., 2009). Orangutans are large brained and have broad behavioral repertoires (van Schaik, 2013). They occupy a complex foraging niche with many food items requiring pre-ingestive processing, including tool use in some populations (van Schaik et al., 1996). Dependent offspring begin to consistently feed on solid food from ∼1 year of age and acquire skill and knowledge repertoires via a mix of individual and social learning, mainly by closely observing their mothers and soliciting food from her, followed by independent practice (Jaeggi et al., 2010; Mikeliban et al., 2021; Schuppli et al., 2016). Because male orangutans start to disperse during adolescence, they have less time to learn from their mothers than their female peers who stay in their natal area and regularly meet with their mothers throughout their lives (Ashbury et al., 2020; Ehmann et al., 2021).

Here, we investigated the effects of mother-offspring characteristics (mother’s parity and offspring age and sex), an ecological factor (prevailing fruit availability), and social factors (association size and the presence of males) on nine maternal behaviors in Sumatran orangutans (*Pongo abelii*). These behaviors fall under four potential domains of maternal investment, which we refer to as maternal functions, namely locomotory support (carrying offspring while moving), skill acquisition support (feeding in close proximity, food transfer), protective proximity maintenance (initiating body contact or close proximity with offspring, following offspring), and independence promotion (terminating body contact or close proximity with offspring, avoiding offspring; here the motivation can either be selfish and/or intended to stimulate independence). We predict that Sumatran orangutan mothers will flexibly adjust their investment in these maternal functions in response to mother-offspring characteristics and socio-ecological factors (given the trade-offs that these factors impose), to ultimately increase their lifetime fitness by possibly optimizing reproductive pace (via the speed at which their offspring develop), offspring survival, and offspring reproduction while ensuring their own survival.

### Predictions

#### 1. Mother’s parity

We expect maternal investment to vary as a function of maternal experience in that primiparous mothers would show more protective proximity maintenance and locomotory and skill acquisition support but less independence promotion toward their offspring than multiparous mothers to compensate for the lack of direct maternal experience.

#### 2. Offspring’s age

As the developing offspring acquires subsistence skills and becomes increasingly competent with age, we expect mothers to show less protective proximity maintenance and provide less locomotory and skill acquisition support on the one hand but increase offspring independence promotion on the other hand.

#### 3. Offspring’s sex

Based on the Trivers-Willard hypothesis, we expect mothers to generally invest more in male than female offspring.

#### 4. Food availability

Due to significant energetic constraints faced by orangutans, we expect maternal investment, especially potentially energy-intensive forms of investment such as locomotory and skill acquisition support (see discussion), to reduce during periods of extremely low food availability.

#### 5. Association size

As a result of the combination of low forest productivity, general energetic constraints, and the energetic requirements of offspring care, we expect associations at certain times (e.g., during large associations) to be costly for females with dependent offspring. We, therefore, expect mothers to reduce their investment in especially potentially energy-intensive forms of investment such as locomotory and skill acquisition support when association sizes are large.

#### 6. Male presence

Because of the potentially high infanticide risk, we expect to see maternal counterstrategies to infanticide by males in that mothers increase protective proximity maintenance and reduce independence promotion in the presence of associating males.

## METHODS

### a. Study site and data collection

For this study, we used data collected on wild Sumatran orangutans from 2007 to 2022 at the Suaq Balimbing research area in the Gunung Leuser National Park (3°42′N, 97°26′E) in South Aceh, Indonesia. Multiple experienced researchers and field assistants collected the data, whose concordance index had reached more than 85% during interobserver reliability testing (computed using simultaneous follow data collected by multiple observers on the same focal individual without any information exchange between the observers). We collected the data through scan sampling (Altmann, 1974) at 2-minute intervals by following mother-offspring pairs, when possible, from their morning to evening nest (see ab.mpg.de/571325/standarddatacollectionrules_suaq_detailed_jan204.pdf for data collection protocols). At each scan, we noted down the behavior of the focal animal, the distance to all association partners, (if distance changed) the individual responsible for a distance change, and whether the offspring was being carried by its mother. We defined associations as individuals staying within 50 m of one another. For the behavior ‘food transfer’ (see below), aside from scan data, we used data collected *ad libitum*. We used a total of 1220 focal follows (including 839 follows on mothers and 381 follows on dependent offspring) for a total duration of 11,271.5 follow hours on 22 mother-offspring pairs (Table S1). We used data on offspring from birth till 8 years of age, as at Suaq, the period of constant mother-offspring association ends at around 8 years, which marks the end of the offspring dependency period (van Noordwijk et al., 2018). As we followed the orangutans opportunistically, depending on encountering them in the study area, sample sizes vary within mother-offspring pairs across behaviors (depending on whether a behavior was recorded during the follow) and between mother-offspring pairs.

### b. Measurement of maternal investment

We calculated maternal investment as the probability that mothers showed a maternal behavior (i.e., proportion of scans during which the focal mother showed a behavior) (Table S2) representing four maternal functions, namely locomotory support, skill acquisition support, protective proximity maintenance, and independence promotion. To minimize the influence of offspring behavior on the expression of the maternal investment, where possible, we accounted for the opportunities that the mothers had available to show the respective behaviors, by including them in the denominator of the calculation (Table S2). The total number of follows and observation hours for the different maternal behaviors included in our analyses are summarized in Figure S1. In orangutans, lactation increases the mother’s energy needs by 25% of baseline level (van Noordwijk et al., 2013). Despite nursing being a significant form of maternal investment in orangutans (van Noordwijk et al., 2013), we did not examine suckling behavior in the current study because in this orangutan population nursing cannot be accurately recorded, as it often happens in day or night nests where visibility of the dependent offspring is low.

### c. Predictors of maternal investment

We modelled the effects of mother’s parity (primiparous/multiparous), offspring age (in years, Figure S2) and sex (female/male), prevailing fruit availability (assessed through a fruit availability index, henceforth called FAI, Figure S3), daily average association size (i.e., the number of association partners apart from the focal individual averaged across the scans taken during a follow, Figure S4), and any adult males in association (presence/absence). FAI was calculated per month as the percentage of fruiting trees in the study site’s two phenology transects that run North-South and West-East (see https://www.aim.uzh.ch/en/research/orangutannetwork/psp.html), comprising approximately 1000 trees. We included maternal parity instead of maternal age as a proxy for maternal experience as a predictor since we did not know the ages of all our adult females.

Dependent offspring were, per definition, in constant association with their mothers. Mothers of different reproductive status (primiparous, multiparous) were followed, with data available on one to three dependent offspring per mother (Table S1). The number of male and female offspring, primiparous and multiparous females, and the number of follows with and without a male varied among the different behaviors. In total, there were 6 primiparous and 16 multiparous mothers and 6 female and 16 male offspring (Table S1). There was at least one adult male associating with the focal mother-offspring pair in ∼42% of the 1220 follows.

### d. Statistical analysis

All statistical analyses were conducted in R (version 4.2.2; R Core Team, 2023). We fitted Generalized Linear Mixed Effects Models (GLMMs; Baayen, 2008) using the *glmer* function of the *lme4* package (version 1.1.34; Bates et al., 2014). Each full model contained maternal investment in a given behavior as a response variable and all the predictors described above as fixed effects. Based on visual inspection of the raw data and the fact that certain behaviors are not expressed at certain ages (e.g., offspring do not feed on solid food when they are very young), we additionally tested for a quadratic effect of offspring age in the models for the probability of mothers feeding in close proximity with their offspring and the probability of food transfer. We did so by comparing the model with only a linear effect of offspring age to a model with both a linear and quadratic effect of offspring age, while keeping all other fixed effects and the random effects the same between the two models, using likelihood ratio tests (LRT; Dobson & Barnett, 2018) as implemented in the *anova* function in R. If the quadratic model outperformed the original full model, the quadratic model was then considered as the full model, else the model with only the linear age effect was considered as the full model. Based on visual inspection, we included only a linear effect of FAI.

Since there were multiple data points for each mother and as certain mothers had data on more than one offspring (*N*=6 of 15 mothers), mother identity was included as a random factor to avoid pseudo-replication, and offspring identity was nested within mother identity to account for potential differences in maternal investment by mothers towards their different offspring, respectively. Since maternal behaviors were calculated in the form of discrete probabilities (numerator and denominator were number of scans, Table S2), we used a logistic model (binomial error distribution and logit link function; McCullagh and Nelder, 1989) to analyze the behaviors and added follow ID as a random factor (we elaborate on the rationale in the supplementary material). To avoid a model being overconfident regarding the precision of fixed effects estimates and to keep the type I error rate at the nominal level of 5%, we included all theoretically identifiable random slopes (Barr et al., 2013; Schielzeth & Forstmeier, 2009). Specifically, we included random slopes of offspring age, FAI, daily average association size, and male presence/absence (the latter manually dummy coded and centered) within mother identity and within offspring identity. We did not include correlation parameters between random slopes and intercepts in our final full models (we elaborate on the rationale in the supplementary material). Prior to fitting the models, we inspected whether all the quantitative predictors were roughly symmetrically distributed. We then z-transformed the quantitative predictors – offspring age, daily average association size, and FAI, for easier interpretable model estimates (Schielzeth, 2010) and to ease model convergence.

After fitting each model, we assessed normality of BLUPs (Best Linear Unbiased Predictors; Harrison et al., 2018; function written by RM), relative model complexity (i.e., the number of observations divided by the number of estimated model parameters), absence of collinearity (using the *vif* function of the library *car* (version 3.1.2; Fox and Weisberg, 2011), maximum Variance Inflation Factor in our models was 1.7; Quinn and Keough, 2002), and overdispersion (maximum dispersion parameter in our models was 0.6; function written by RM) for all the models. After fitting the full model, we tested the fit of the full model against the null model (Forstmeier & Schielzeth, 2011), which contained only the random effects, using LRT as implemented in the *anova* function in R. If the full model produced a significantly better fit than the null model, we tested the significance of the individual fixed effects by using the *drop1* function, which drops one fixed effect at a time and compares the fit of the resulting model to that of the full model. We used a cut-off of *P* < 0.05 for significance. We report the odds ratio (i.e., the exponent of the estimate) for significant fixed effects, which is a measure of effect size in logistic regression (Breaugh, 2003). For a given fixed effect, an odds ratio of <1 indicates a reduction in maternal investment, while values >1 indicate an increase in maternal investment. We further report marginal coefficients of determination (i.e., the variance explained by the fixed effects; R^2^m) and conditional coefficients of determination (i.e., the variance explained by the entire model; R^2^c) obtained using the delta method of the *r.squaredGLMM* function of the MuMin package (version 1.47.5; Barton and Barton, 2015). To assess the uncertainty in the estimated individual effects, we obtained 95% confidence intervals using bootstrapping (*N*=1000 bootstraps; function written by RM). To avoid unreliable estimation of the variance caused by the included random factors, we included only those offspring with data from a minimum of five calendar days (i.e., five focal follows).

#### Ethical Note

The research protocol of this study was approved by the Indonesian State Ministry for Research, Technology and Higher Education (BRIN RISTEK; Research Permit No.: 152/SIP/FRP/SM/V/2012) and complied with the legal requirements of Indonesia. This study was purely observational and complied with the Principles for the Ethical Treatment of Nonhuman Primates by the American Society of Primatologists (2001).

#### Data Availability

The datasets analyzed during the current study and the source codes for all the analyses and graphs presented in the main text and supplementary material of the manuscript are available in the Harvard Dataverse repository, https://doi.org/10.7910/DVN/8WV8XX.

## RESULTS

Extent of maternal investment varied from behavior to behavior (Figure S5). For the behavior ‘feeding in close proximity’, the model with the linear and quadratic age effects performed significantly better (considered subsequently as the full model) than the model with only the linear age effect (*P*<0.001). For the behavior ‘food transfer’, the model with only the linear age effect performed equally well (considered subsequently as the full model) as a model with both the linear and quadratic age effects (*P*=0.479). The loglikelihoods of final full models without correlation parameters between random intercepts and slopes did not differ much from the models with the correlation parameters (Table S3). The full model explained significantly more variation in maternal investment than the corresponding null model for six of the nine maternal behaviors (Table S4). Below we elaborate on the results of the individual fixed effects on these six behaviors. The full models for the behaviors ‘follow’, ‘avoid’, and ‘food transfer’ were not significantly different from their respective null models (Table S4).

### 1. Mother’s parity

Parity did not significantly predict maternal investment in any of the maternal behaviors (Table 1).

**Table 1.** Results of the supported full models investigating six maternal behaviors of wild Sumatran orangutans (*Pongo abelii*) as a function of mother-offspring characteristics and socioecological factors using data collected between 2007 and 2022 at the Suaq Balimbing research area in South Aceh, Indonesia. FAI stands for prevailing fruit availability index; P/A stands for presence/absence; *R^2^m* stands for the marginal coefficient of determination; *R^2^c* stands for the conditional coefficient of determination. Significant *P* values (*P*<0.05) are marked in bold.

| Term | Estimate | SE | Lower CI | Upper CI | Z value | P value |
| --- | --- | --- | --- | --- | --- | --- |
| *Probability of mother initiating body contact with her offspring ($R^2_m$: 0.57; $R^2_c$: 0.91)* |  |  |  |  |  |  |
| Intercept | -7.077 | 0.270 | -7.617 | -6.522 | † | † |
| Mother's parity (Primiparous) | -0.363 | 0.312 | -0.924 | 0.266 | -1.163 | 0.240 |
| Offspring age* | -1.715 | 0.196 | -2.069 | -1.319 | -8.757 | **<0.001** |
| Offspring sex (Male) | 0.481 | 0.262 | -0.048 | 1.023 | 1.831 | 0.062 |
| FAI* | 0.096 | 0.135 | -0.121 | 0.303 | 0.709 | 0.510 |
| Average association size* | 0.223 | 0.086 | 0.018 | 0.342 | 2.597 | **0.022** |
| Male P/A (P) | 0.212 | 0.257 | -0.169 | 0.659 | 0.823 | 0.401 |
| *Probability of mother terminating body contact with her offspring ($R^2_m$: 0.34; $R^2_c$: 0.94)* |  |  |  |  |  |  |
| Intercept | -2.847 | 0.366 | -3.609 | -2.068 | † | † |
| Mother's parity (Primiparous) | 0.811 | 0.454 | -0.128 | 1.744 | 1.783 | 0.082 |
| Offspring age* | 1.034 | 0.284 | 0.495 | 1.544 | 3.643 | **0.004** |
| Offspring sex (Male) | 0.310 | 0.402 | -0.519 | 1.040 | 0.773 | 0.452 |
| FAI* | 0.006 | 0.057 | -0.109 | 0.106 | 0.099 | 0.923 |
| Average association size* | -0.088 | 0.064 | -0.266 | 0.032 | -1.374 | 0.182 |
| Male P/A (P) | -0.206 | 0.128 | -0.472 | 0.046 | -1.607 | 0.121 |
| *Probability of mother initiating close proximity with her offspring ($R^2_m$: 0.15; $R^2_c$: 0.65)* |  |  |  |  |  |  |
| Intercept | -4.187 | 0.306 | -4.837 | -3.583 | † | † |
| Mother's parity (Primiparous) | -0.318 | 0.304 | -0.941 | 0.306 | -1.046 | 0.283 |
| Offspring age* | -0.484 | 0.065 | -0.644 | -0.341 | -7.414 | **0.026** |
| Offspring sex (Male) | -0.008 | 0.287 | -0.698 | 0.644 | -0.028 | 0.978 |
| FAI* | 0.021 | 0.080 | -0.184 | 0.151 | 0.259 | 0.803 |
| Average association size* | -0.002 | 0.129 | -0.214 | 0.174 | -0.016 | 0.988 |
| Male P/A (P) | 0.224 | 0.124 | -0.014 | 0.492 | 1.801 | 0.095 |
| *Probability of mother terminating close proximity with her offspring ($R^2_m$: 0.38; $R^2_c$: 0.90)* |  |  |  |  |  |  |
| Intercept | -3.769 | 0.332 | -4.380 | -2.984 | † | † |
| Mother's parity (Primiparous) | -0.707 | 0.367 | -1.446 | -0.027 | -1.928 | 0.071 |
| Offspring age* | 0.885 | 0.174 | 0.564 | 1.217 | 5.100 | **<0.001** |
| Offspring sex (Male) | -0.337 | 0.345 | -1.188 | 0.288 | -0.977 | 0.336 |
| FAI* | -0.123 | 0.090 | -0.274 | 0.041 | -1.377 | 0.143 |
| Average association size* | -0.137 | 0.061 | -0.266 | -0.020 | -2.241 | **0.024** |
| Male P/A (P) | -0.043 | 0.114 | -0.302 | 0.180 | -0.381 | 0.743 |
| *Probability of mother feeding in close proximity with her offspring ($R^2_m$: 0.51; $R^2_c$: 0.99)* |  |  |  |  |  |  |
| Intercept | -5.016 | 1.004 | -6.750 | -2.642 | † | † |
| Mother's parity (Primiparous) | 2.100 | 1.010 | 0.292 | 4.010 | 2.079 | 0.056 |
| Offspring age* | 2.406 | 0.133 | 2.094 | 2.706 | 18.139 | <**0.001** |
| (Offspring age)^2* | -1.812 | 0.323 | -2.422 | -1.283 | -5.603 | <**0.001** |
| Offspring sex (Male) | 2.773 | 1.045 | 1.026 | 4.513 | 2.652 | **0.008** |
| FAI* | -0.419 | 0.160 | -0.748 | -0.093 | -2.613 | **0.014** |
| Average association size* | -0.088 | 0.132 | -0.404 | 0.134 | -0.663 | 0.504 |
| Male P/A (P) | 0.121 | 0.197 | -0.223 | 0.525 | 0.611 | 0.612 |
| *Probability of mother carrying her offspring (R2m: 0.74; R2c: 0.98)* |  |  |  |  |  |  |
| Intercept | -2.866 | 0.960 | -4.959 | -0.927 | † | † |
| Mother's parity (Primiparous) | -1.109 | 1.028 | -3.443 | 0.817 | -1.079 | 0.304 |
| Offspring age* | -4.615 | 0.388 | -5.660 | -3.949 | -11.890 | <**0.001** |
| Offspring sex (Male) | 0.723 | 0.984 | -1.424 | 2.908 | 0.735 | 0.460 |
| FAI* | 0.456 | 0.341 | -0.128 | 0.957 | 1.338 | 0.174 |
| Average association size* | -0.039 | 0.091 | -0.190 | 0.164 | -0.422 | 0.662 |
| Male P/A (P) | 0.496 | 0.207 | 0.122 | 0.970 | 2.397 | **0.025** |

†Not shown due to limited interpretability. \*z-transformed to a mean of 0 and a SD of 1.
Original mean and SD are reported in the supplementary material (Table S5).

### 2. Offspring age

We found that offspring age significantly predicted maternal investment in all six maternal behaviors (Table 1). Whereas the probability of mothers initiating body contact and close proximity with their offspring and carrying their offspring decreased with offspring age (odds ratio for an increase in offspring age: 0.18, 0.62, 0.01, respectively; Figure 1a, 1b, 1f), the probability of mothers terminating body contact and terminating close proximity increased with offspring age (odds ratio: 2.81, 2.42, respectively; Figure 1c, 1d). The probability of mothers feeding in close proximity with their offspring had a quadratic relationship with offspring age, with the probability being initially low, then increasing with offspring age; subsequently, it reached a peak between five and six years of age and decreased thereafter (odds ratio: linear age-effect: 11.09; quadratic age-effect: 0.16; Figure 1e).

**Figure 1.**
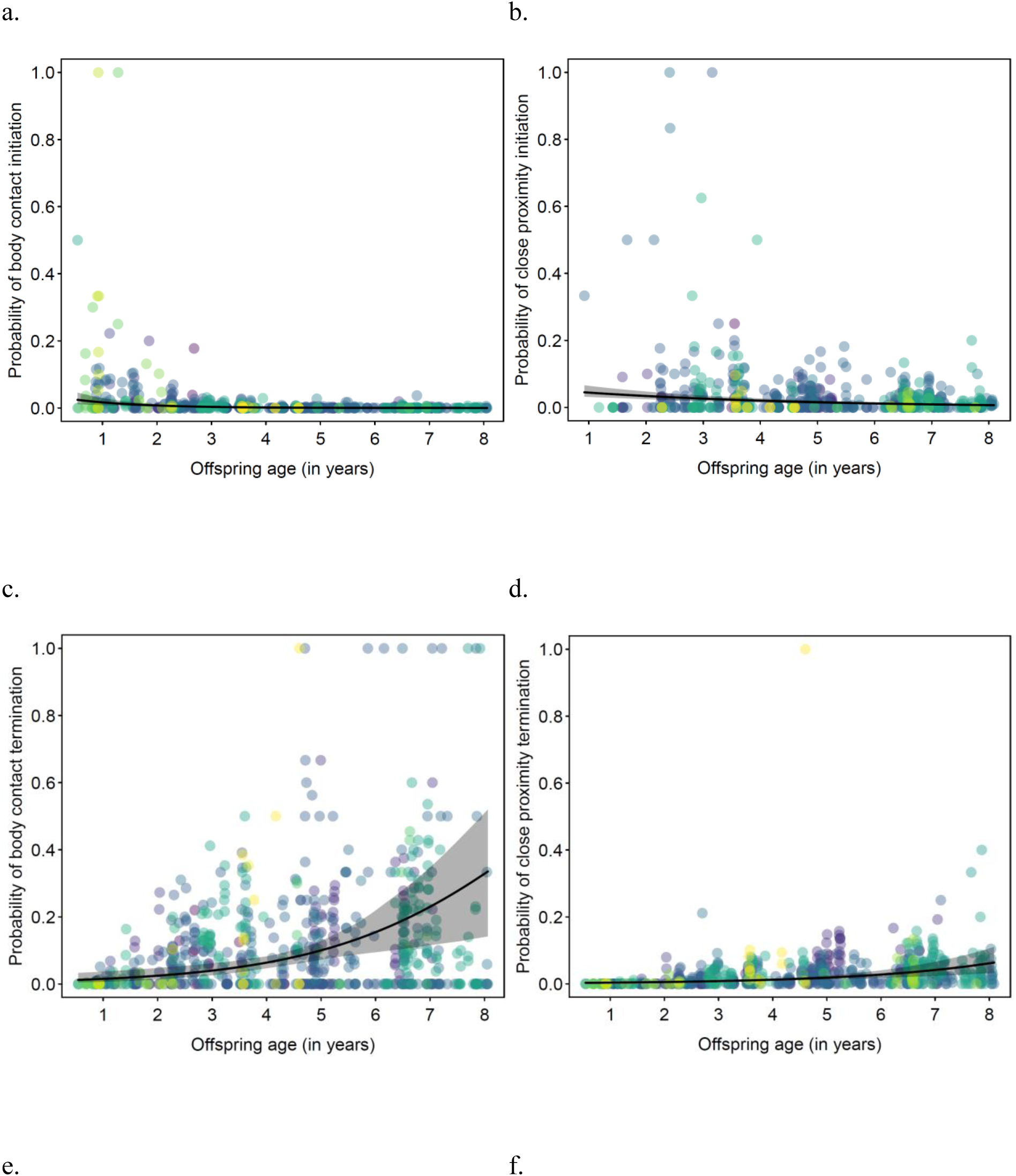
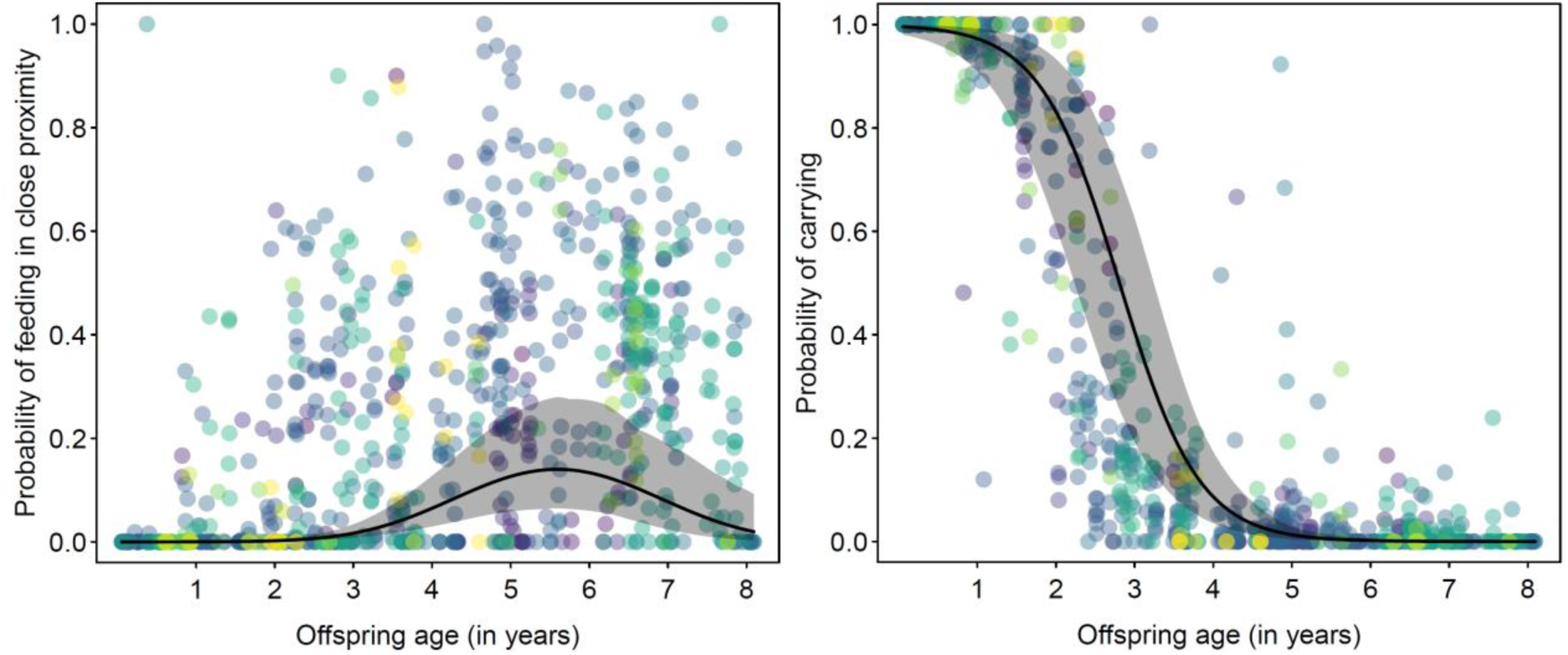
Probability of mothers a) initiating body contact and b) initiating close proximity, c) terminating body contact, d) terminating close proximity, e) feeding in close proximity, and f) carrying their offspring as a function of their offspring’s age in wild Sumatran orangutans (*Pongo abelii*) analyzed using data collected between 2007 and 2022 at the Suaq Balimbing research area in South Aceh, Indonesia. Each dot represents a follow, and they are colored according to offspring identity (color for each offspring is shown in supplementary material, Figure S1). The solid black line represents the fitted model (offspring age, prevailing fruit availability index, and association size are at their average; mother’s parity, offspring sex, and male presence/absence were manually dummy coded and then centered). The shaded area in grey represents the 95% CI of the fitted model.

### 3. Offspring sex

We found that the probability of mothers feeding in close proximity with their offspring was significantly higher for mothers with a male offspring than for those with a female offspring (odds ratio: 16.0; Table 1; Figure 2). Offspring sex was not a significant predictor of any of the other maternal behaviors tested (Table 1).

**Figure 2.**
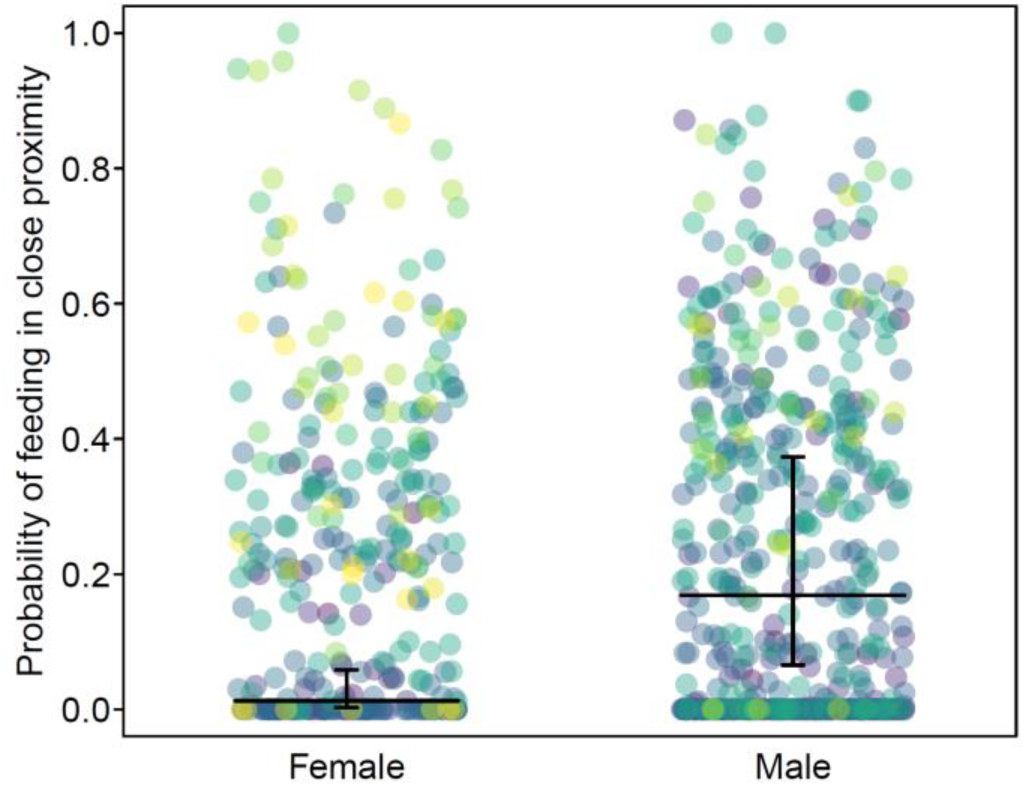
Probability of mothers feeding in close proximity with their offspring as a function of offspring’s sex in wild Sumatran orangutans (*Pongo abelii*) analyzed using data collected between 2007 and 2022 at the Suaq Balimbing research area in South Aceh, Indonesia. Each dot represents a follow, and they are colored according to offspring identity (color for each offspring is shown in supplementary material, Figure S1). The horizontal black lines represent the fitted model (offspring age, prevailing fruit availability index, and association size are at their average; mother’s parity and male presence/absence were manually dummy coded and then centered). The vertical black lines represent the 95% CI of the fitted model. The data points are jittered on the x-axis to reduce overlap.

### 4. FAI

During the study period, FAI ranged between 3.37 and 16.71, with a mean ± SD = 10.035 ± 2.875 (Figure S3). We found that with an increase in FAI there was a significant reduction in the probability of a mother feeding in close proximity with her offspring (odds ratio: 0.66; Table 1; Figure 3), FAI was not a significant predictor of any of the other maternal behaviors tested (Table 1).

**Figure 3.**
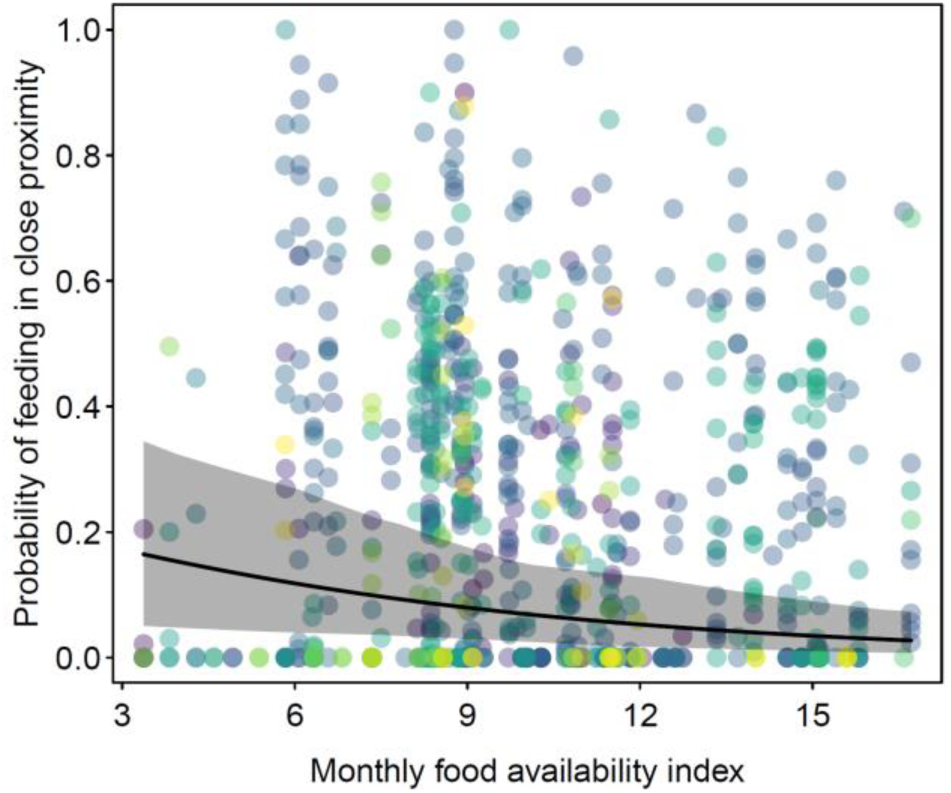
Probability of mothers feeding in close proximity with their offspring as a function of monthly prevailing fruit availability index in wild Sumatran orangutans (*Pongo abelii*) analyzed using data collected between 2007 and 2022 at the Suaq Balimbing research area in South Aceh, Indonesia. Each dot represents a follow, and they are colored according to offspring identity (color for each offspring is shown in supplementary material, Figure S1). The solid black line represents the fitted model (offspring age, prevailing fruit availability index, and association size are at their average; mother’s parity, offspring sex, and male presence/absence were manually dummy coded and then centered). The shaded grey area represents the 95% CI of the fitted model.

### 5. Average association size

During the study period, the daily average number of association partners for the focal mother or offspring ranged between 0.89 (excluding the focal individual) to 11.75 association partners (mean ± SD = 1.655 ± 0.823; averaged across all follows). We found that with an increase in average association size, there was a significant increase in the probability of the mother initiating body contact (odds ratio: 1.25; Table 1; Figure 4a) and a significant decrease in the probability of mothers terminating close proximity (odds ratio: 0.87; Table 1; Figure 4b) with her offspring. Average association size did not significantly predict any of the other maternal behaviors tested (Table 1).

**Figure 4.**
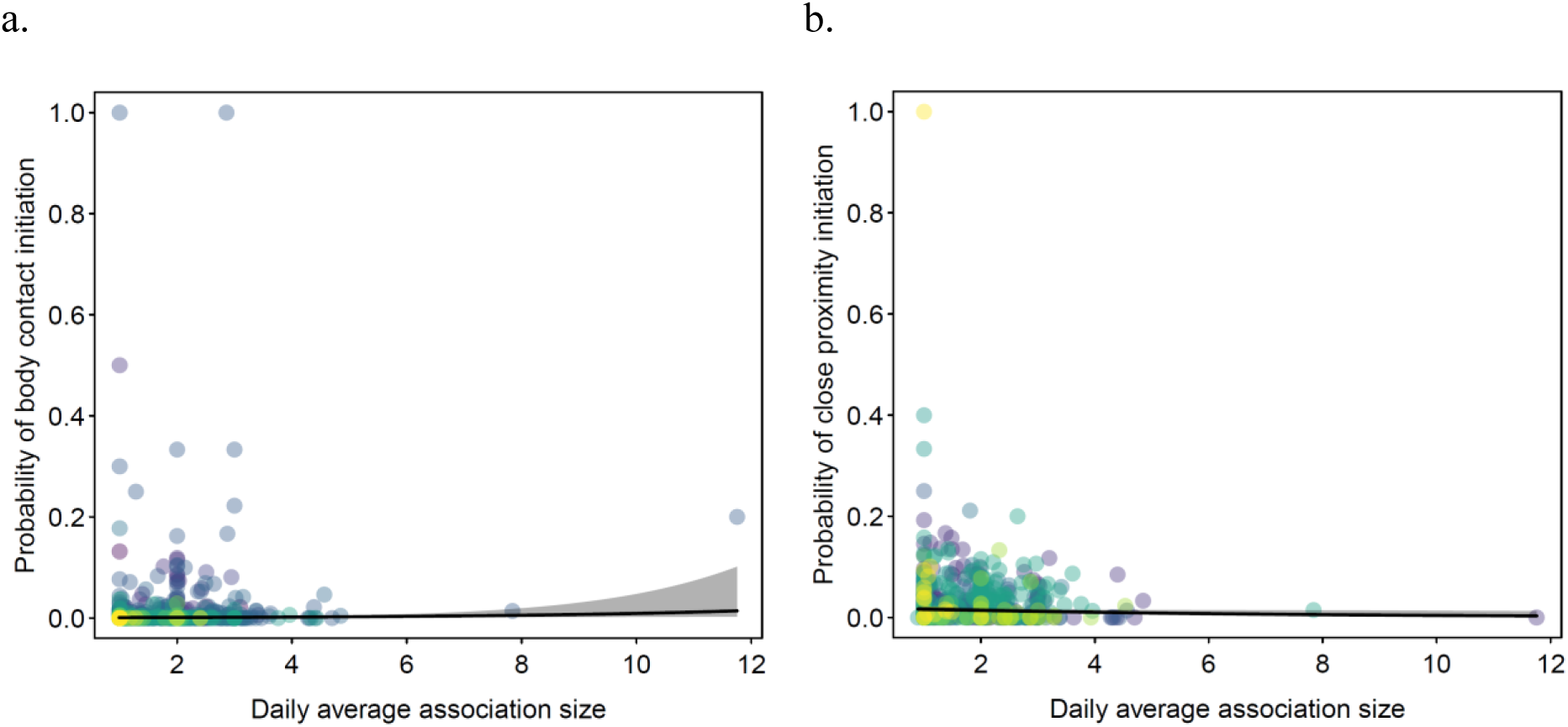
Probability of mothers a) initiating body contact and b) terminating close proximity with their offspring as a function of their daily average association size in wild Sumatran orangutans (*Pongo abelii*) analyzed using data collected between 2007 and 2022 at the Suaq Balimbing research area in South Aceh, Indonesia. Each dot represents a follow, and they are colored according to offspring’s identity (color for each offspring is shown in supplementary material, Figure S1). The solid black line represents the fitted model (offspring age, prevailing fruit availability index, and average association size are at their average; mother’s parity, offspring sex, and male presence/absence were manually dummy coded and then centered). The shaded grey area represents the 95% CI of the fitted model.

### 6. Male presence

Male presence in associations had a significant positive effect on the probability of mothers carrying their offspring while moving (odds ratio: 1.64; Table 1; Figure5). Male presence/absence did not significantly predict any of the other maternal behaviors (Table 1).

**Figure 5.**
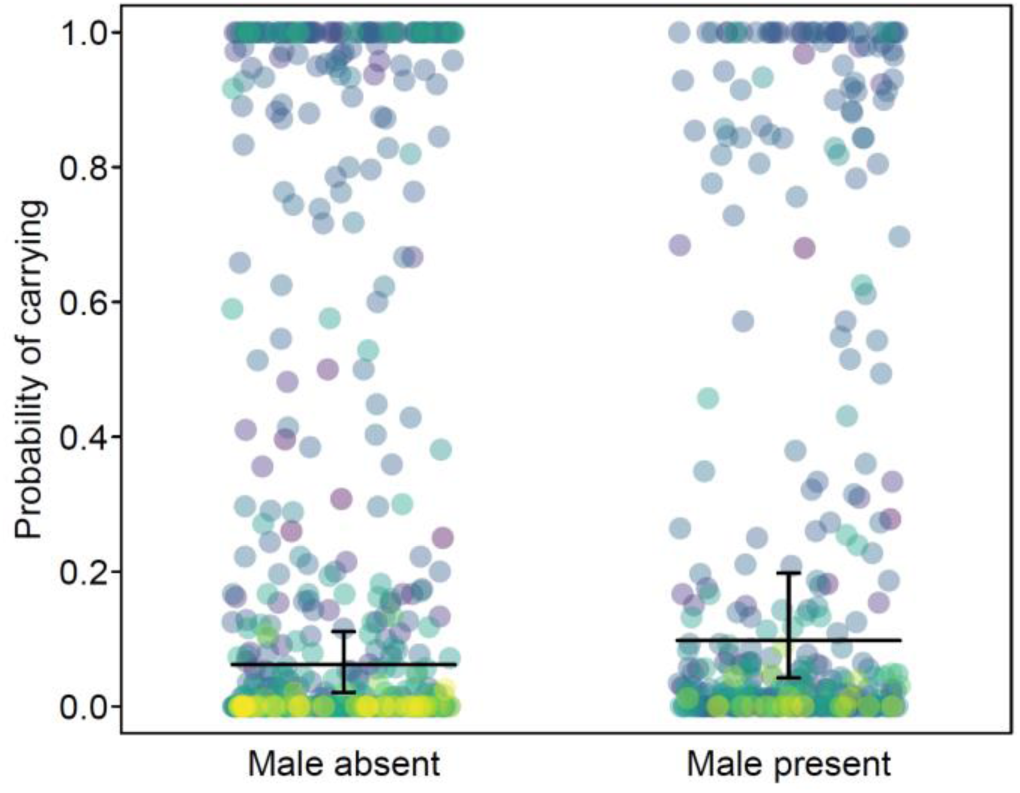
Probability of mothers carrying their offspring as a function of male presence/absence in associations in wild Sumatran orangutans (*Pongo abelii*) analyzed using data collected between 2007 and 2022 at the Suaq Balimbing research area in South Aceh, Indonesia. Each dot represents a follow, and they are colored according to offspring identity (color for each offspring is shown in supplementary material, Figure S1). The horizontal black lines represent the fitted model (offspring age, prevailing fruit availability index, and association size are at their average; mother’s parity and offspring’s sex were manually dummy coded and then centered). The vertical black lines represent the 95% CI of the fitted model. The data points are jittered on the x-axis to reduce overlap.

## DISCUSSION

Our study showed that mother-offspring characteristics and socioecological variables are associated with variation in maternal investment in several maternal behaviors in Sumatran orangutans. However, in the current study, we did not investigate the extent to which the analyzed maternal behaviors entail actual costs (e.g., energetic and/or opportunity costs) for the mother, and thus represent forms of maternal investment in the classical sense (Clutton-Brock, 1991). Below we discuss these results from a fitness perspective and in light of what is known about maternal investment in primates.

### Mother parity

Contrary to our first prediction, primiparous and multiparous mothers did not significantly differ in their maternal investment in locomotory and skill acquisition support, protective proximity maintenance, and independence promotion. Our results augment the existing literature showing a lack of difference in maternal investment based on parity in terrestrial primates (cradle, restrain, retrieve: Seay, 1966; different spatial proximity measures: Silk, 1991). However, these findings contrast with studies on primates of varying degree of terrestriality in which multiparous mothers seem to invest more into (premasticated) food transfer (e.g., chimpanzee (*Pan troglodytes*): Bădescu et al., 2020) and independence promotion (e.g., rhesus macaques: Seay, 1966; Japanese macaques (*Macaca fuscata*): Tanaka, 1989) during the first six months of (offspring) age. With a larger sample size, follow up studies should investigate the interaction effect of mother’s parity and offspring age on maternal investment.

The fact that first time and experienced mothers were similar in their investment through different maternal functions suggests that primiparous Sumatran orangutan females do not need to compensate for a lack of experience and show adequate offspring age-appropriate care without prior direct experience. Given the mounting evidence for social learning of routine skills by immature orangutans (Schuppli et al., 2016; Schuppli and van Schaik 2019), it is plausible that daughters learn maternal behaviors from their mothers either during their own dependency period (Maestripieri, 2005) or by watching their mothers’ behavior towards their younger sibling (Berman, 1990). Analyzing developmental speed and survival of offspring of mothers of differing experience would help to pin down whether primiparous mothers are indeed fully competent caregivers.

### Offspring age

Offspring age affected six of the maternal behaviors. In support of our second prediction, mothers decreased locomotory and skill acquisition support and protective proximity maintenance, while they increased independence promotion as their dependents became older. This pattern suggests that mothers adjust their investment to their offspring’s needs, and thus optimize their investment in terms of energy and time costs without negatively affecting their offspring’s development. Examples of maternal investment changing with offspring age in terrestrial and arboreal primates include contact termination (Rhesus monkeys: Berman, 1980) increasing with offspring age, and time spent carrying (Mantled howler monkeys (*Alouatta palliata*): Dias et al., 2018) and body contact initiation (Rhesus monkeys: Berman, 1980) decreasing with offspring age.

The initial low probability of mothers and offspring feeding in close proximity to their (while they fed on the same food item) is a result of the offspring starting to feed on solid foods only gradually from around 1 year of age onwards (van Noordwijk et al., 2013; van Noordwijk & van Schaik, 2005). This was followed by an increase in the probability of mothers and offspring feeding in close proximity, peaking around 5-6 years of offspring age, which coincides with a steep increase in the immatures’ dietary breadth (Schuppli et al., 2021). Previous chimpanzee and orangutan studies also showed that mothers were increasingly less likely to share food items, as food processing competence of the offspring increases with age (e.g., Bădescu et al., 2020; Jaeggi et al., 2008; Mikeliban et al., 2021), indicating that mothers may be attuned to the developing competence of their offspring (Mikeliban et al., 2021). However, our current analysis cannot disentangle the role of mother vs. offspring in the probability of feeding in close proximity. With a larger dataset, one could analyze whether the mother or the offspring breaks close proximity while feeding. Such an analysis would help to understand if maternal contribution to feeding skill acquisition changes with offspring age.

### Offspring sex

In line with our expectation, we found that mothers with male offspring showed higher skill acquisition support through feeding in close proximity than those with female offspring. Compared to their female age-peers, immature males peer at their mothers less frequently (Ehmann et al., 2021). They may thus need more skill acquisition support from their mothers (see also results on chimpanzees: Estienne et al., 2019; Lonsdorf, 2006). Research on another orangutan population has shown that females reach larger diet repertoires sooner than males (Schuppli et al., 2021). Due to earlier skill acquisition, female offspring can afford to feed further away from their mothers, suggesting that this behavior may be initiated by the offspring more than that by the mothers.

Mothers investing more into male offspring appears to be in line with the Trivers-Willard hypothesis (Trivers and Willard, 1973), given that orangutan males have a higher potential reproductive outcome than females. However, we did not account for maternal body condition and our sample size did not allow comparisons of the same mothers with offspring of both sexes. Therefore, support for the Trivers-Willard hypothesis (Trivers and Willard, 1973) by our results is very limited. Furthermore, our analyses cannot rule out that the difference in skill acquisition support are not mostly mediated by the offspring: male and female offspring may differ in how much time they spend at different distances to their mothers, which may lead to differences in close proximity feeding.

Contrary to our third prediction, mothers with male or female offspring did not differ in their investment in protective proximity maintenance, locomotory and other forms of skill acquisition support, and independence promotion. These results are in line with other studies on primates which found little to no overt sex-biased difference in maternal investment in behaviors related to protective proximity maintenance and independence promotion (restraint, rejection, body contact, and various proximity measures: Brown and Dixson, 2000; Silk, 1991; Tanaka, 1989). The absence of sex-biased maternal investment in these behaviors could be a result of these behaviors being vital for the development and safety of both the sexes. It is possible that sex-biased investment occurs during a certain age period rather than throughout offspring development. Our sample size did not allow us to test the interaction between offspring age and sex on maternal investment. Furthermore, we had more than twice the number of male than female offspring in our dataset, which means that our current results, including the absence of sex differences in certain behaviors, should be treated with caution. A larger and a more balanced sample would allow us to interpret the effect of offspring sex more reliably.

### Food availability

Contrary to our fourth prediction, locomotory support did not decrease with decreasing food availability. Since our measures of skill acquisition support included either solid food sharing or food patch sharing (in case of feeding in close proximity) with one’s offspring, we expected such support to affect mother’s feeding efficiency, and thus ultimately result in lower energy intake for her. However, contrary to our prediction, skill acquisition support through mothers feeding in close proximity to their offspring increased with decreasing food availability. These patterns suggest that food availability may not constrain the time or energy that mothers invest in locomotory or skill acquisition support, however we had only few data points in the low range of FAI, and hence the results should be interpreted with caution. At times of low fruit availability, orangutans feed on fallback foods such as bark, pith, and leaves (Vogel et al., 2017). Even though food availability is generally high at Suaq in comparison to other orangutan habitats (Marshall et al., 2009), there is an increase in bark and pith feeding during periods of relatively low food availability in Suaq (Schuppli, unpublished data). Some of these fallback foods are more clumped (e.g., bark), and may thus lead to higher frequency of feeding in close proximity than while feeding on fruits (which at Suaq are often eaten in fruiting trees with large crowns). Furthermore, some of the fallback foods are difficult to process (e.g., bark and pith), i.e., require more intense learning (Schuppli et al., 2016), and may thus favor increased time spent in close proximity.

We further found that food availability did not affect any other maternal behavior. A reason for this could be that food availability is generally high in Suaq. In other populations where individuals face more pronounced fluctuations, punctuated with periods of extremely low food availability (Marshall et al., 2009), maternal investment may be considerably affected by prevailing food availability.

All in all, the results regarding the effects of food availability on maternal investment add to the growing body of literature that shows that resource availability influences certain aspects of maternal care. It is well known that in primates food availability affects maternal body condition (e.g., Dias et al., 2018) and that food availability (Hauser, 1993) and maternal body condition influence the quality of maternal care (Clutton-Brock, 1991; Fairbanks & McGuire, 1995; Lee et al., 1991). Mothers living in high quality habitat and who are in better body condition are more likely to respond to their offspring’s distress calls (Hauser, 1993) and remain spatially closer to their offspring or carry their offspring (Dias et al., 2018) than those living in poor quality habitat or who are in poor body condition.

### Association size

Contrary to our fifth prediction, energy-intensive forms of maternal behavior, such as locomotory and skill acquisition support, were not affected by daily average association sizes. However, in agreement with studies on terrestrial primates (e.g., Berman et al., 1997), protective proximity maintenance with dependents through body contact initiation increased and close proximity termination decreased with an increase in daily average association size, even though notably the effect sizes were small. Average association size did not influence any of the other maternal behaviors. Even though Suaq orangutans are more socially tolerant than Bornean orangutans (*Pongo pygmaeus wurmbii*), they are semi-solitary with low overall rates of close associations and social interactions, most likely because associations come with energetic costs such as increased duration of the active period and decreased feeding time (Kunz et al., 2021). The presence of association partners may be a safety risk for young and socially inexperienced offspring, thereby prompting an increase in protective proximity maintenance from their mothers. This indicates that the costs that mothers are experiencing through these associations are not constraining their maternal investment in these functions or at least not their immediate investment (during the time of the associations). However, since we did not have dense data on maternal investment during large association sizes, these results should be interpreted with caution. A previous study on orangutans found a positive relationship between association size and offspring-initiated mother-offspring conflict (Falkner, 2015). In combination with the findings of the current study, this suggests that mothers are more eager to keep their dependent offspring close to them during associations than their dependents are. Future studies should investigate how the effects of association size on maternal behavior depend on the age-sex class or the degree of relatedness/familiarity of the associating individuals, with data spanning the entire range of the predictor.

### Male presence

In support of our sixth prediction, we found that mothers were more likely to carry their offspring in the presence than in the absence of an associating male. In several terrestrial and arboreal primate species with sexual size dimorphism, mothers were reported to show potential anti-infanticide strategies such as increased body contact time, initiation of body contact, physical restraint, as well as inspection of offspring, and reduced distance and decreased time away from their offspring in the presence of males (vervets: Fairbanks & McGuire, 1987; chimpanzees: Otali and Gilchrist, 2006; Bornean orangutans: Scott et al., 2023, 2019). Because of their arboreal lifestyle, carrying is likely an additional means by which orangutan mothers protect their offspring in the presence of a male and keep the risk of falling low for offspring. Our results indicate that Sumatran orangutan females may view at least some males as a potential threat and increase locomotory support as a counter-infanticide strategy. Mothers might also change their behavior in the presence of males depending on their reproductive status, which is closely linked to the age of their current dependent offspring (Fairbanks & McGuire, 1987; Kunz et al., 2022; Scott et al., 2019). Furthermore, it is likely that not all males are a threat, and maternal behavior may vary based on whether a male is a resident or not (Fairbanks & McGuire, 1987; Kunz et al., 2022; van Noordwijk et al., 2023). With a larger sample size, future studies should test the interaction effect of offspring age and male presence/absence and the effect of associating male’s familiarity on maternal investment.

### Unaffected maternal behaviors

The probabilities of mothers following and avoiding their offspring and the probability of food transfer were not significantly associated with any of the six predictors that we tested. Previous studies have shown that mothers adjust how much they tolerate food taking by their offspring according to the age of their offspring and based on the properties of the food item that is taken (Jaeggi et al., 2008; Mikeliban et al., 2021). However, the sample for food transfer was quite low in the current study, especially in relation to the complexity of our statistical model, which may be the reason for the full model not performing better than the null model. With respect to the behaviors ‘follow’ and ‘avoid’, it remains to be investigated whether they do indeed represent forms of maternal investment in orangutans. Notably, these were among the most infrequent behaviors in our data.

## CONCLUSION

Our comprehensive approach of testing the effects of six predictors on nine maternal behaviors showed that Sumatran orangutan mothers invest in locomotory and skill acquisition support, protective proximity maintenance, and independence promotion flexibly in response to mother-offspring characteristics and short-term varying socioecological factors. Offspring age, sex, association size, and male presence had similar effects on maternal investment in Sumatran orangutans as found in terrestrial primates, underscoring the importance of these shared effects across species and the environments they inhabit. In many aspects, these flexible adjustments appear to support offspring development or protect the mothers from facing costs that may endanger their own survival. As such, this study is in line with the evolutionary theory that females allocate investment in a flexible manner to maximize their lifetime reproductive success (Clutton-brock, 1991, Trivers, 1972, Lee et al., 1991). However, whether maternal investment through these behaviors indeed translates into actual fitness benefits for the mother (through increased own or offspring survival, and/or faster reproductive rates) remains to be investigated.

### Inclusion and Diversity Statement

The author list includes a contributor from the location where the research was conducted, who participated in interpretation of the findings and long-term project administration.

## Acknowledgements

We thank all the dedicated field staff, students, and research assistants (in particular, Armas Fitra, Adami Khairul, Fikar Zulkarnean, Julia Kunz, Lara Nellissen, Mudin, Natasha Bartolotta, Saidi Agan, Toni, and Ulil Azhari) who contributed to the data collected for this study at Suaq. We are grateful to the Indonesian State Ministry for Research and Technology (BRIN-RISTEK), the Directorate General of Natural Resources and Ecosystem Conservation-Ministry of Environment and Forestry of Indonesia (KSDAE-KLHK), the Ministry of Internal Affairs, Indonesia, the Sumatran orangutan conservation Program (SOCP), and Balai Besar Taman Nasional Gunung Leuser (BBTNGL) in Medan for their permission and support to conduct this study. We thank Universitas Nasional (UNAS) for support and collaboration. We thank the Max Planck Institute of Animal Behavior, University of Zurich, A.H. Schultz Foundation, Leakey Foundation (Primate Research Fund and project grant), SUAQ Foundation, and the Volkswagen Stiftung (Freigeist fellowship to CS) for financial support. Open access funding was provided by the Max Planck Institute of Animal Behavior.

## Contributions

Conceptualization: TR, MvN, CS.

Data curation: TR, CS.

Formal analysis: TR, RM, PB, CS.

Funding acquisition: CS.

Investigation: TR, RM, SSUA, PB, MvN, CS.

Field project administration: SSUA, CS.

Supervision: RM, CS.

Writing—original draft: TR.

Writing—review & editing: TR, RM, SSUA, PB, MvN, CS.

