## Supplemental Material for "Maternal behavior in Sumatran orangutans (*Pongo abelii*) is modulated by mother-offspring characteristics and socioecological factors"

*Glossary*

**Maternal behavior**: Any behavior shown by a mother towards her offspring after birth.

**Maternal care**: Any presumably offspring fitness-increasing behavior shown by a mother after birth.

**Maternal investment**: Reproductive effort put forth by a mother for her current developing offspring, which reduces her future fecundity.

**METHODS**

Table S1. Overview of the focal individuals of wild Sumatran orangutans (*Pongo abelii*) that were studied between 2007 and 2022 at the Suaq Balimbing research area in South Aceh, Indonesia, with at least 5 focal follows.

| **SI.**  **No.** | **Mother ID** | **Offspring**  **ID** | **Mother's**  **parity** | **Offspring's**  **estimated DOB** | **Offspring's**  **sex** | **Sampling age period (min, max years)** |
| --- | --- | --- | --- | --- | --- | --- |
| 1 | Alice | Amor | Multiparous | 01-Jan-15 | Male | 0.13-5.14 |
| 2 | Chick | Chuck | Multiparous | 01-Jul-07 | Male | 0.55-0.82 |
| 3 | Cissy | Chindy | Multiparous | 01-Jan-03 | Female | 5.03-7.85 |
| 4 | Cissy | Cinnamon | Multiparous | 01-Apr-12 | Female | 0.11-7.04 |
| 5 | Dodi | Dalia | Multiparous | 01-Oct-13 | Female | 0.13-1.04 |
| 6 | Dodi | Diddy | Primiparous | 01-Dec-05 | Male | 3.20-5.82 |
| 7 | Ellie | Eden | Primiparous | 01-Nov-14 | Female | 0.06-7.22 |
| 8 | Friska | Frankie | Multiparous | 01-Aug-12 | Male | 0.80-7.96 |
| 9 | Friska | Fredy | Multiparous | 01-Jun-05 | Male | 2.09-8.05 |
| 10 | Intai | Inky | Multiparous | 01-Jul-03 | Male | 4.86-7.87 |
| 11 | Lilly | Luther | Primiparous | 01-Mar-16 | Male | 0.57-3.94 |
| 12 | Lisa | Leon | Multiparous | 15-Nov-18 | Male | 0.05-0.52 |
| 13 | Lisa | Lilly | Primiparous | 01-Mar-01 | Female | 6.59-7.92 |
| 14 | Lisa | Lois | Multiparous | 01-Aug-10 | Male | 0.52-7.85 |
| 15 | Nora | Nuk | Multiparous | 01-Jan-06 | Male | 3.10-5.27 |
| 16 | Piniata | Pepito | Multiparous | 01-Jan-13 | Male | 4.56-6.95 |
| 17 | Raffi | Rendang | Multiparous | 15-Jul-13 | Male | 0.57-1.28 |
| 18 | Raffi | Ronaldo | Multiparous | 01-Jan-06 | Male | 1.66-5.63 |
| 19 | Sarabi | Sazu | Primiparous | 01-Jul-06 | Male | 7.28-7.62 |
| 20 | Sarabi | Simba | Multiparous | 01-Mar-13 | Male | 0.61-2.09 |
| 21 | Sonya | Sound | Primiparous | 01-Aug-18 | Female | 3.05-3.15 |
| 22 | Tiara | Tornado | Multiparous | 01-Jul-14 | Male | 2.18-5.11 |

| a.  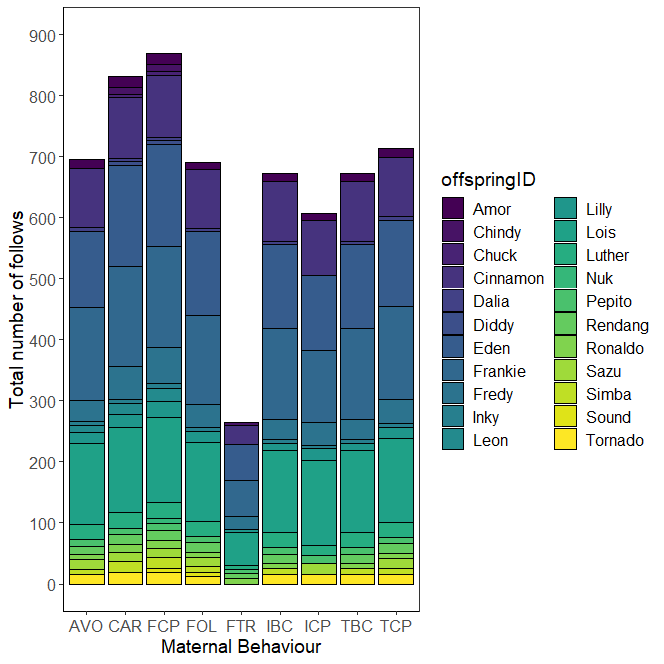 | b.  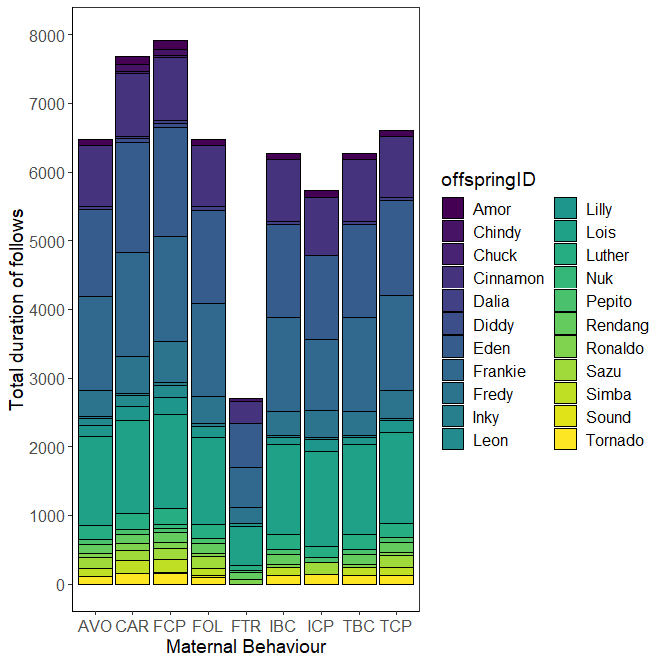 |
| --- | --- |

Figure S1. Plot of the total a.) number of follows and b.) duration of follows (hours) for each offspring with at least 5 follows for each maternal behavior in wild Sumatran orangutans (*Pongo abelii*). Data were collected between 2007 and 2022 at the Suaq Balimbing research area in South Aceh, Indonesia. AVO: avoid; CAR: carry; FCP: feed in close proximity; FOL: follow; FTR: food transfer; IBC: initiate body contact; ICP: initiate close proximity; TBC: terminate body contact; TCP: terminate close proximity.

Table S2. Maternal behaviors in wild Sumatran orangutans (*Pongo abelii*) observed between 2007 and 2022 at the Suaq Balimbing research area in South Aceh, Indonesia, including the numerator and denominator used to calculate maternal investment while considering the opportunities that mothers had available to show each behavior. The behaviors fall under four maternal functions: ^(1)^Locomotory support, ^(2)^Skill acquisition support, ^(3)^Protective proximity maintenance, and ^(4)^Independence promotion.

| **Behavior** | **Numerator** | **Denominator** |
| --- | --- | --- |
| ^(1)^Carry | No. of scans in which the mother carried her offspring | Total no. of scans in which the mother moved during a follow |
| ^(2)^Feeding in close proximity* | No. of scans in which the mother was feeding within 5 m of her offspring who was also feeding on the same food item as its mother | Total no. of scans during which the mother was feeding during a follow |
| ^(2)^Food transfer† | No. of offspring begging events with food taken from mother by offspring | No. of times offspring was begging from its mother during a follow |
| ^(3)^Initiate body contact | No. of scans in which the mother initiated body contact with her offspring (>0 m to 0 m) | Total no. of scans in which the mother and her offspring were not in contact (i.e., >0 m) during a follow |
| ^(3)^Initiate close proximity | No. of scans in which the mother initiated close proximity with her offspring (>5-<=50 m to <=5 m but no body contact) | Total no. of scans in which the mother and offspring were not in close proximity (i.e., >5-<=50 m) during a follow |
| ^(3)^Follow | No. of scans in which the mother moved closer to her offspring immediately (i.e., within 2-min period) after her offspring moved away from her | Total no. of scans in which the offspring moved away (to any distance) from its mother during a follow |
| ^(4)^Avoid | No. of scans in which the mother moved away from her offspring immediately (i.e., within 2-min period) after her offspring moved closer to her | Total no. of scans in which the offspring moved closer to its mother during a follow |
| ^(4)^Terminate body contact | No. of scans in which the mother had terminated body contact with her offspring (0 m to >0 m) | Total no. of scans in which the mother and offspring were in body contact during a follow |
| ^(4)^Terminate close proximity | No. of scans in which the mother terminated close proximity with her offspring (<=5 m, but no contact to >5-<=50 m) | Total no. of scans in the which mother and offspring were in close proximity (i.e., <=5 m, but no contact) during a follow |

*Mother feeding in close proximity with her dependent offspring on the same food item (i.e., providing opportunities for observational and interactive social learning in the feeding context; a food item is defined as a combination of a species and a plant part that is eaten, following Bastian et al., 2010)

†Food transfer during offspring begging events (begging is defined as “any attempt by offspring to obtain food from its mother”; Jaeggi et al., 2008). >95% of the food transfer events involve mothers tolerating offspring’s taking of food from them.

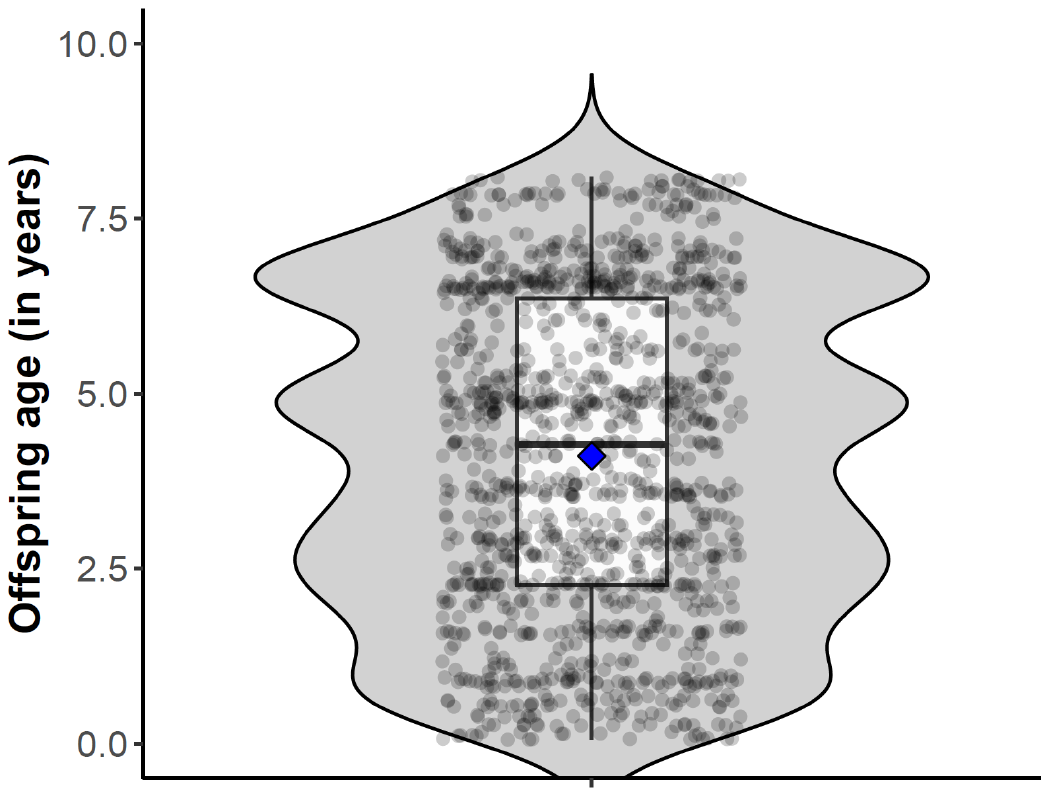

Figure S2. Distribution of offspring age (in years) over all follows on wild Sumatran orangutans (*Pongo abelii*) collected between 2007 and 2022 at the Suaq Balimbing research area in South Aceh, Indonesia. Each dot represents offspring age during one follow. The data points are jittered along the x-axis to reduce their overlap. The width of the violin plot represents the density of data points at a specific point. The boxplot shows the lower quartile, median, mean (in blue diamond), and upper quartile, along with the whiskers.

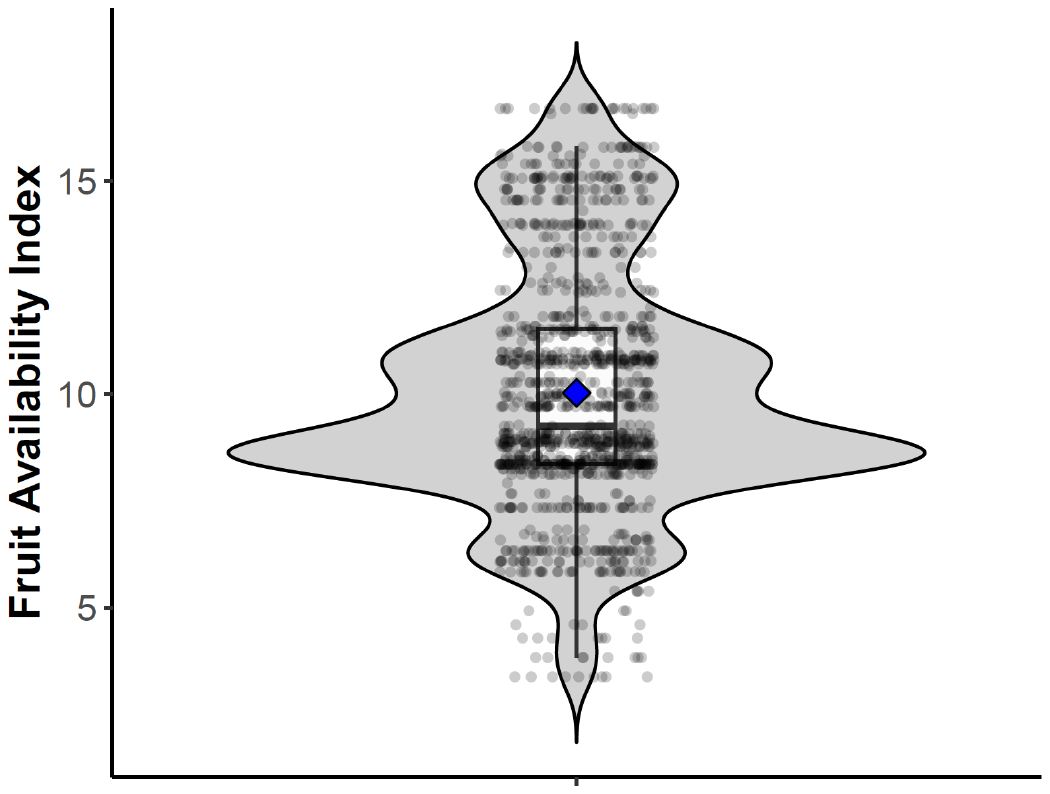

Figure S3. Distribution of fruit availability index (FAI) over all the follows of wild Sumatran orangutans (*Pongo abelii*) collected between 2007 and 2022 at the Suaq Balimbing research area in South Aceh, Indonesia. Each dot represents the FAI during one follow. The data points are jittered along the x-axis to reduce their overlap. The width of the violin plot represents the density of data points at a specific point. The boxplot shows the lower quartile, median, mean (in blue diamond), and upper quartile, along with the whiskers.

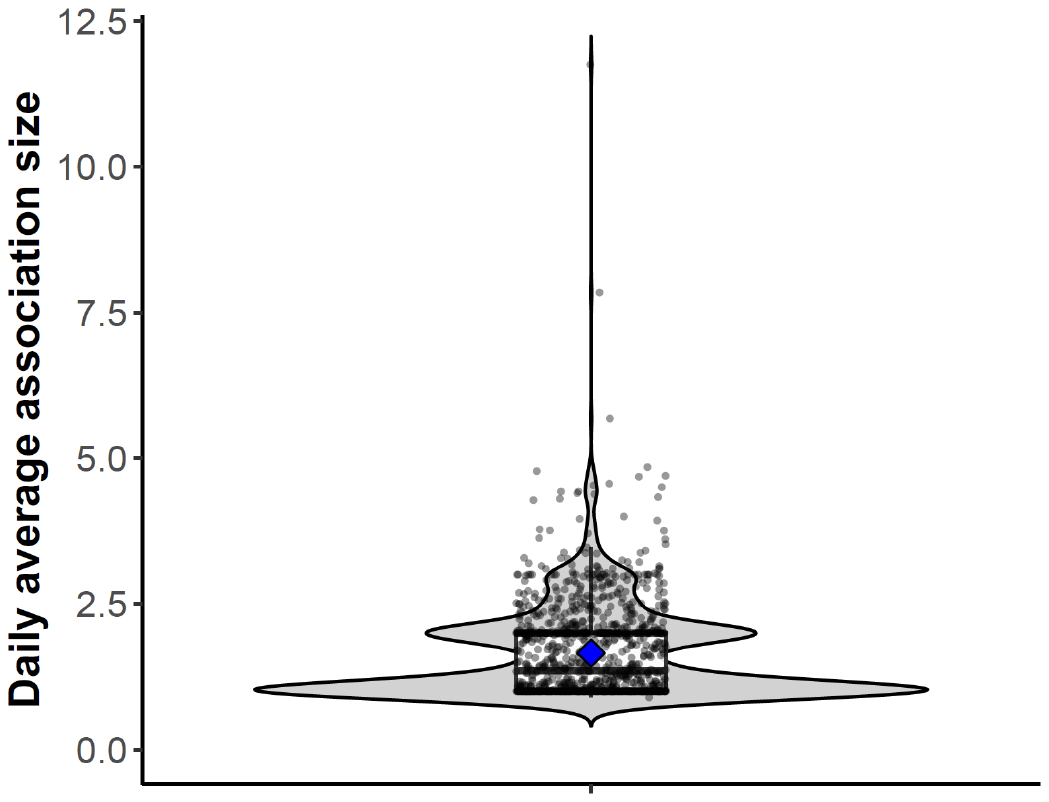

Figure S4. Distribution of daily average association size over all the follows of wild Sumatran orangutans (*Pongo abelii*) collected between 2007 and 2022 at the Suaq Balimbing research area in South Aceh, Indonesia. Each circle represents the average association size (averaged across the scans) during one follow. The data points are jittered along the x-axis to reduce their overlap. The width of the violin plot represents the density of data points at a specific point. The boxplot shows the lower quartile, median, mean (in blue diamond), and upper quartile, along with the whiskers.

*Analysis of discrete probabilities in R*

In R, discrete probabilities can be analyzed using binomial models with the response variable handed over as a matrix with two columns, in which the first column denotes the number of scans in which the mother showed a behavior, and the second column denotes the number of scans in which the mother did not show a behavior (Baayen, 2008). When the binary response matrix is handed over to the *glmer* function, it is analyzed on a trial-by-trial basis (i.e., *glmer* models individual outcomes and not the proportions per follow). Hence, to account for the observations in the same row belonging to the same follow, a random intercept effect that has one level for each row in the response matrix (i.e., random effect of follow ID) was included in the model.

*Maximal model structure: correlation between random intercepts and random slopes*

Originally, we began with a maximal model (Barr et al., 2013), including parameters for the correlations among random intercepts and slopes. However, correlation parameters were usually estimated to have absolute values being essentially one, which is indicative of them not being identifiable (Matuschek et al., 2017). Hence, we removed the correlation parameters, which did not lead to a large reduction in model fit (log likelihoods with and without the correlation parameters are shown in Table S5).

**RESULTS**

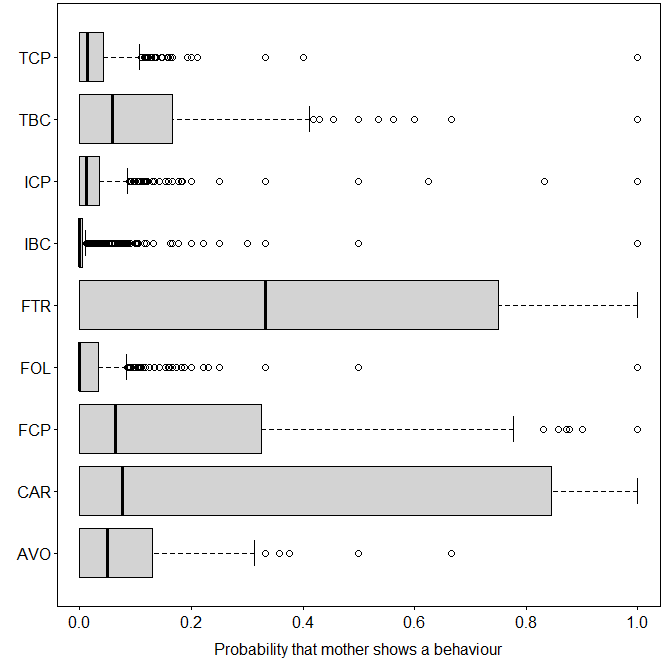

Figure S5. Boxplot of maternal investment (i.e., the probability of mother showing a behavior given the opportunities available to mothers to show that behavior) in the nine maternal behaviors in wild Sumatran orangutans (*Pongo abelii*) analyzed using the data collected between 2007 and 2022 at the Suaq Balimbing research area in South Aceh, Indonesia. The boxplot shows minimum, first quartile, median, third quartile, and maximum for each behavior. Whiskers and individual data points outside the whiskers are also shown. AVO: avoid; CAR: carry; FCP: feed in close proximity; FOL: follow; FTR: food transfer; IBC: initiate body contact; ICP: initiate close proximity; TBC: terminate body contact; TCP: terminate close proximity.

Table S3. Comparison of log-likelihoods between models with and without correlation parameters (1,2, respectively) between random intercept and random slopes within mother identity and offspring identity for nine maternal behaviors in wild Sumatran orangutans (*Pongo abelii*) analyzed using the data collected between 2007 and 2022 at the Suaq Balimbing research area in South Aceh, Indonesia.

| **Model** | **df** | **Loglik** |
| --- | --- | --- |
| *Probability of mother initiating body contact with her offspring* | | |
| 1 | 38 | -712.61 |
| 2 | 18 | -714.58 |
| *Probability of mother terminating body contact with her offspring* | | |
| 1 | 38 | -1323.11 |
| 2 | 18 | -1328.37 |
| *Probability of mother initiating close proximity with her offspring* | | |
| 1 | 38 | -962.28 |
| 2 | 18 | -970.25 |
| *Probability of mother terminating close proximity with her offspring* | | |
| 1 | 38 | -1530.31 |
| 2 | 18 | -1537.80 |
| *Probability of mother following her offspring* | | |
| 1 | 38 | -479.95 |
| 2 | 18 | -482.65 |
| *Probability of mother avoiding her offspring* | | |
| 1 | 38 | -1021.83 |
| 2 | 18 | -1024.18 |
| *Probability of food transfer* | |  |
| 1 | 38 | -307.07 |
| 2 | 18 | -312.41 |
| *Probability of mother feeding in close proximity with her offspring* | | |
| 1 | 51 | -1454.24 |
| 2 | 21 | -1464.88 |
| *Probability of mother carrying her offspring* | | |
| 1 | 38 | -3134.46 |
| 2 | 18 | -3146.31 |

Table S4. Full-null model comparison for nine maternal behaviors in wild Sumatran orangutans (*Pongo abelii*) analyzed using the data collected between 2007 and 2022 at the Suaq Balimbing research area in South Aceh, Indonesia. The full models contained all the fixed and random effects. The null models contained only the random intercept and slope effects. Significant *P* values (*P*<0.05) are marked in bold. Model complexity (i.e., the number of observations per estimated parameter) of the full model is shown next to each maternal behavior.

|  | **Chisq** | ***df*** | ***P*** |
| --- | --- | --- | --- |
| *Probability of mother initiating body contact with her offspring (model complexity: 37.4)* | | | |
| Full model | 41.094 | 6 | **<0.001** |
| *Probability of mother terminating body contact with her offspring (model complexity: 37.4)* | | | |
| Full model | 17.724 | 6 | **0.007** |
| *Probability of mother initiating close proximity with her offspring (model complexity: 33.7)* | | | |
| Full model | 15.280 | 6 | **0.018** |
| *Probability of mother terminating close proximity with her offspring (model complexity: 39.7)* | | | |
| Full model | 26.327 | 6 | **0.000** |
| *Probability of mother following her offspring (model complexity: 38.4)* | | | |
| Full model | 5.402 | 6 | 0.493 |
| *Probability of mother avoiding her offspring (model complexity: 38.6)* | | | |
| Full model | 3.925 | 6 | 0.687 |
| *Probability of food transfer (model complexity: 14.7)* | | | |
| Full model* | 12.432 | 6 | 0.053 |
| *Probability of mother feeding in close proximity with her offspring (model complexity: 41.4)* | | | |
| Full model† | 65.857 | 7 | **<0.001** |
| *Probability of mother carrying her offspring (model complexity: 46.2)* | | | |
| Full model | 40.597 | 6 | **<0.001** |

* Model with only a linear effect of offspring age; adding a quadratic effect of offspring age did not improve model fit.

†Model with a linear and quadratic effect of offspring age; adding a quadratic effect of offspring age improved model fit.

Table S5. Original means and standard deviations of the z-transformed quantitative predictors for the maternal behaviors in wild Sumatran orangutans (*Pongo abelii*) analyzed using the data collected between 2007 and 2022 at the Suaq Balimbing research area in South Aceh, Indonesia. These behaviors significantly differed from their respective null models. FAI stands for prevailing food availability index.

| **Predictor** | **Mean (SD)** |
| --- | --- |
| *Probability of mother initiating body contact with her offspring* | |
| Offspring age | 4.312 (2.096) |
| FAI | 10.195 (2.81) |
| Average association size | 1.683 (0.883) |
| *Probability of mother terminating body contact with her offspring* | |
| Offspring age | 4.312 (2.096) |
| FAI | 10.195 (2.81) |
| Average association size | 1.683 (0.883) |
| *Probability of mother initiating close proximity with her offspring* | |
| Offspring age | 5.144 (1.799) |
| FAI | 10.269 (2.798) |
| Average association size | 1.637 (0.79) |
| *Probability of mother terminating close proximity with her offspring* | |
| Offspring age | 4.46 (2.133) |
| FAI | 10.165 (2.816) |
| Average association size | 1.701 (0.884) |
| *Probability of mother feeding in close proximity with her offspring* | |
| Offspring age | 4.053 (2.322) |
| FAI | 10.125 (2.861) |
| Average association size | 1.677 (0.836) |
| *Probability of mother carrying her offspring* | |
| Offspring age | 4.077 (2.334) |
| FAI | 10.066 (2.794) |
| Average association size | 1.679 (0.837) |
